# Human CXCL17 Activates and Binds to Fish GPR25 Orthologs

**DOI:** 10.1101/2025.06.10.658772

**Authors:** Wen-Feng Hu, Juan-Juan Wang, Jie Yu, Jun-Jie Yao, Ya-Li Liu, Zeng-Guang Xu, Zhan-Yun Guo

## Abstract

C-X-C motif chemokine ligand 17 (CXCL17) functions as a chemoattractant, though its receptor has been controversial. Recent independent studies, including our own, identified CXCL17 as an agonist for the orphan G protein-coupled receptor 25 (GPR25). While GPR25 orthologs are found across fishes to mammals, CXCL17 orthologs appear to be mammalian-specific, leaving the endogenous ligand for non-mammalian GPR25 orthologs unknown. This study unexpectedly found that human CXCL17 exhibits high activity towards GPR25 orthologs from the zebrafish (*Danio rerio*) and coelacanth (*Latimeria chalumnae*). Recombinant human CXCL17 efficiently activated both fish GPR25 orthologs in a NanoLuc Binary Technology (NanoBiT)-based β-arrestin recruitment assay, and induced chemotactic movement in transfected human embryonic kidney (HEK) 293T cells expressing fish GPR25. A human CXCL17 mutant lacking three C-terminal residues showed no such effect. A NanoBiT-based binding assay revealed that a SmBiT-tagged human CXCL17 C-terminal fragment specifically bound to secretory large NanoLuc fragment (sLgBiT)-fused fish GPR25 orthologs. Fish GPR25 orthologs had significantly higher cell surface expression in transfected HEK293T cells compared to human GPR25, improving β-arrestin recruitment assay data quality. Despite approximately 400 million years of divergence between humans and fishes, the high activity of human CXCL17 on fish GPR25 orthologs suggests that the CXCL17–GPR25 pair may be conserved across all vertebrates, even though non-mammalian CXCL17 orthologs remain unidentified.

## Introduction

C-X-C motif chemokine ligand 17 (CXCL17) was identified in 2006 [1,2] and acts as a chemoattractant for various immune cells, including T cells, monocytes, macrophages, and dendritic cells [1,3–15]. It is also implicated in tumor development, likely by regulating tumor immunity [2,16–21]. The functions of CXCL17 are presumed to be mediated by specific plasma membrane receptor(s). Initially, CXCL17 was reported as an agonist of the orphan G protein-coupled receptor 35 (GPR35) in 2015 [22], but subsequent studies did not support this interaction [23–25]. More recently, CXCL17 has been described as an agonist of the orphan MAS-related receptor MRGPRX2, an allosteric inhibitor of the chemokine receptor CXCR4, or a chemokine signaling modulator through binding to extracellular glycosaminoglycans [25–27]. Most recently, Ocón’s group and our group independently identified CXCL17 as an agonist for the orphan G protein-coupled receptor 25 (GPR25) [28,29], a rarely studied A-class G protein-coupled receptor (GPCR) primarily expressed in some immune cells, such as T cells, plasma cells, and B-cells, according to data from the Human Protein Atlas (https://www.proteinatlas.org).

According to the National Center for Biotechnology Information (NCBI) gene database, GPR25 orthologs are broadly distributed from fishes to mammals, whereas CXCL17 orthologs appear to be exclusive to extant mammals. This disparity renders the endogenous ligand of non-mammalian GPR25 orthologs elusive. A previous study reported that apelin and apela, two peptide hormones highly conserved across all vertebrates, act as weak agonists for some non-mammalian GPR25 orthologs [30]. GPR25 shares low sequence similarity with the apelin receptor (APLNR), suggesting that apelin and apela likely exhibit low cross-activity with non-mammalian GPR25 orthologs rather than serving as their true endogenous agonists.

In our recent work, functional assays for human GPR25 were challenging because of its low cell surface expression in transfected human embryonic kidney (HEK) 293T cells [29]. The previous report indicated that zebrafish (*Danio rerio*) GPR25 (herein designated as Dr-GPR25) has a higher cell surface expression level than human GPR25, according to the microscopic fluorescence images of the transfected HEK293 cells expressing the enhanced green fluorescent protein-tagged zebrafish or human GPR25. Given this, we hypothesized that human CXCL17 might possess cross-activity with fish GPR25 orthologs, and that utilizing fish GPR25 orthologs could improve assay sensitivity due to their higher cell surface expression level in transfected HEK293T cells.

Therefore, the objective of this study was to investigate the interaction between human CXCL17 and two representative fish GPR25 orthologs: Dr-GPR25 from the ray-finned fish *Danio rerio* (zebrafish) and Lc-GPR25 from the lobe-finned fish *Latimeria chalumnae* (coelacanth). Our results demonstrated that human CXCL17 is highly active to fish GPR25 orthologs despite divergence of humans and fishes approximately 400 million years ago, suggesting that the CXCL17-GPR25 pair might be conserved across all vertebrates, even though CXCL17 orthologs have not yet been identified from non-mammalian vertebrates.

## Materials and Methods

### Preparation of recombinant human CXCL17 proteins

The N-terminally 6×His-tagged wild-type (WT) human CXCL17 and the C-terminally truncated [desC3]CXCL17 were prepared according to our previous procedure [29]. Briefly, the 6×His-tagged WT or mutant CXCL17s were overexpressed in *Escherichia coli* as inclusion bodies, purified via a Ni^2+^ column after solubilization, refolded *in vitro*, and purified via high-performance liquid chromatography (HPLC) using a C_8_ reverse-phase column (Zorbax 300SB-C8, 9.4 × 250 mm, Agilent Technologies, Santa Clara, CA, USA). After lyophilization, the refolded 6×His-tagged WT or mutant CXCL17s were dissolved in 1.0 mM aqueous hydrochloride (pH 3.0), quantified by ultra-violet absorbance at 280 nm using the extinction coefficient of ε_280 nm_ = 11000 M^-1^ cm^-1^, and used for activity assays.

To generate the expression construct for N-terminally 6×His-SmBiT-fused human CXCL17, the encoding DNA fragment was amplified via polymerase chain reaction (PCR) using our previous construct pET/6×His-CXCL17 as template [29] and then ligated into a pET vector via Gibson assembly, resulting in the expression construct pET/6×His-SmBiT-CXCL17 (Fig. S1). Thereafter, the recombinant 6×His-SmBiT-CXCL17 protein was prepared according to the procedure developed for preparation of 6×His-CXCL17 [29].

To generate the expression construct for N-terminally 6×His-SmBiT-fused C-terminal fragment (26 residues) of human CXCL17, a DNA fragment encoding 6×His-SmBiT-CXCL17-C26 was chemically synthesized and a DNA fragment encoding the *Mycobacterium xenopi*-derived intein was PCR amplified using our previous construct pET/FAM237A-intein-6×His as template [31]. Thereafter, the two DNA fragments were ligated into a pET vector via Gibson assembly, resulting in the expression construct pET/6×His-SmBiT-CXCL17-C26-intein (Fig. S1). The intein fusion protein was overexpressed in the *E. coli* strain BL21(DE3) as inclusion bodies according to standard protocol. After solubilization from inclusion bodies via an *S*-sulfonation approach, the intein fusion protein was purified via a Ni^2+^ column, treated with 50 mM dithiothreitol at room temperature for 30 min, cooled on ice and subjected to 20-fold dilution by slow addition of ice-cold 20 mM phosphate buffer (pH7.4). After overnight incubation at room temperature, the mixture was loaded onto a sulfonic propyl (SP) ion-exchange column, and the released 6×His-SmBiT-CXCL17-C26 was eluted from the column by 1.0 M NaCl solution, and further purified by HPLC using a C_18_ reverse-phase column (Zorbax 300SB-C18, 9.4 × 250 mm, Agilent Technologies). After lyophilization, the peptide powder of 6×His-SmBiT-CXCL17-C26 was dissolved in 1.0 mM aqueous hydrochloride (pH 3.0), quantified by ultra-violet absorbance at 280 nm using the extinction coefficient of ε_280 nm_ = 1490 M^-1^ cm^-1^, and used for receptor activation or binding assays.

### Preparation of synthetic human apelin and apela

Human apelin-13 and human apela-21 were chemically synthesized at GL Biochem (Shanghai, China) via standard solid-phase peptide synthesis. The crude linear peptides were purified by HPLC via a C_18_ reverse-phase column (Zorbax 300SB-C18, 9.4 × 250 mm, Agilent Technologies) and confirmed by molecular mass. The synthetic linear apela-21 was further subjected to oxidative refolding in 0.5 L-arginine solution (pH7.5) containing 300 μM linear peptide and 900 μM 2,2’-dithiodipyridine (CAS no. 2127-03-9). After incubation on ice for 2 h, the refolding mixture was acidified to pH3-4 and purified by HPLC via a C_18_ reverse-phase column (Zorbax 300SB-C18, 9.4 × 250 mm, Agilent Technologies). The lyophilized peptide powder of the synthetic apelin-13 and apela-21 (refolded) were weighed via a precise balance, dissolved in 1.0 mM aqueous hydrochloride (pH 3.0), and used for activity assays.

### Generation of the expression constructs for fish GPR25 orthologs

The information about the fish GPR25 orthologs was retrieved from the gene database of NCBI and listed in Table S1. The coding region of Dr-GPR25 was PCR amplified using zebrafish genomic DNA as template and synthetic oligoes as primers (Table S2). The PCR product was subjected to electrophoresis and the amplified DNA fragment (∼1.2 kb) was recovered from the agarose gel and ligated into a β-arrestin recruitment assay vector via Gibson assembly (Table S2). The resultant expression construct pTRE3G-BI/Dr-GPR25-LgBiT:SmBiT-ARRB2 coexpresses a C-terminally large NanoLuc fragment (LgBiT)-fused Dr-GPR25 and an N-terminally SmBiT-fused human β-arrestin 2 (ARRB2) under control of a doxycycline (Dox)-response bidirectional promoter. Thereafter, the coding region of Dr-GPR25 in this construct was PCR amplified using appropriate primers and ligated into appropriate vectors as listed in Table S2. The construct PB-TRE/Dr-GPR25 encodes an untagged Dr-GPR25 and the construct PB-TRE/sLgBiT-Dr-GPR25 encodes an N-terminally secretory LgBiT (sLgBiT)-fused Dr-GPR25 under control of a Dox-response promoter.

The coding region of Lc-GPR25 was chemically synthesized at Tsingke Biotechnology (Beijing, China) and ligated into a pcDNA3.1 vector (Table S2). Thereafter, the coding region of Lc-GPR25 in the construct pcDNA3.1/Lc-GPR25 was PCR amplified using appropriate primers and ligated into appropriate vectors as listed in Table S2. The nucleotide sequence of Dr-GPR25 and Lc-GPR25 in these constructs was confirmed by DNA sequencing.

### Generation of a coexpression construct for human TPST1 and TPST2

The expression constructs for human tyrosylprotein sulfotransferase TPST1 and TPST2 in pENTER vector were purchased from WZ Bio (Jinan, Shandong Province, China). Thereafter, their coding regions were PCR amplified using appropriate primers and sequentially ligated into the multiple cloning site 1 (MCS1) and multiple cloning site 2 (MCS2) of the pTRE3G-BI vector (ClonTech, Mountain View, CA, USA) via Gibson assembly. The expression construct pTRE3G-BI/TPST1:TPST2 coexpresses human TPST1 and human TPST2 under control of Dox-response bidirectional promoter. The nucleotide sequence of TPST1 and TPST2 in this construct was confirmed by DNA sequencing.

### The NanoBiT-based **β**-arrestin recruitment assays

The NanoBiT-based β-arrestin recruitment assay was conducted according to our previous procedure developed for Hs-GPR25 and other GPCRs [29,31]. Briefly, HEK293T cells were transiently transfected with a β-arrestin assay construct (pTRE3G-BI/Dr-GPR25-LgBiT:SmBiT-ARRB1, pTRE3G-BI/Lc-GPR25-LgBiT:SmBiT-ARRB2, or pTRE3G-BI/Hs-APLNR-LgBiT:SmBiT-ARRB2) together with the expression control vector pCMV-TRE3G (Clontech) using the transfection reagent Lipo8000 (Beyotime Technology, Shanghai, China). Next day, the transfected cells were trypsinized, seeded into white opaque 96-well plates, and cultured in induction medium (complete DMEM plus 1.0 ng/mL of Dox) at 37 °C for ∼24 h to ∼90% confluence. To start the β-arrestin recruitment assay, the induction medium was removed and pre-warmed activation solution (serum-free DMEM plus 1% BSA) containing NanoLuc substrate was added (40 μL/well, containing 0.5 μL of NanoLuc substrate stock from Promega, Madison, WI, USA), and bioluminescence data were immediately collected for ∼4 min on a SpectraMax iD3 plate reader (Molecular Devices, Sunnyvale, CA, USA) with an interval of 11 s. Subsequently, indicated peptide (diluted in the activation solution) was added (10 μL/well), and bioluminescence data were continuously collected for ∼10 min with an interval of 11 s. The measured absolute bioluminescence signals were corrected for inter well variability by forcing all curves after addition of NanoLuc substrate (without ligand) to same level and plotted using the SigmaPlot 10.0 software (SYSTAT software, Chicago, IL, USA). To obtain the dose curve for CXCL17 activating Dr-GPR25 or Lc-GPR25, the measured bioluminescence data at 583 s were plotted with CXCL17 concentrations using the SigmaPlot 10.0 software.

### Transwell chemotaxis assay

The transwell chemotaxis assay was conducted according to our previous procedure developed for Hs-GPR25 [29]. Briefly, HEK293T cells were transfected with the construct PB-TRE/Dr-GPR25 or PB-TRE/Lc-GPR25 using the transfection reagent Lipo8000 (Beyotime Technology). Next day, the transfected cells were changed to the induction medium (complete DMEM plus 3.0 ng/mL of Dox) and continuously cultured for ∼24 h. Thereafter, the cells were trypsinized, suspended in serum-free DMEM at the density of ∼5×10^5^ cells/mL, and seeded into polyethylene terephthalate membrane (8 μm pore size)-coated permeable transwell inserts (150 μL/well) that were put into the lower plate chambers containing 600 μL of chemotactic agent (serum-free DMEM plus 1% BSA and indicated concentrations of WT or mutant human CXCL17). After cultured at 37 °C for ∼6.5 h, solution in the inserts were removed and cells on the upper face of the permeable membrane were wiped off using cotton swaps, and cells on the lower face of the permeable membrane were fixed with 4% paraformaldehyde solution, stained with crystal violet staining solution (Beyotime Technology), and observed under an Olympus APX100 microscope (Tokyo, Japan).

### The NanoBiT-based ligand-receptor binding assay

The NanoBiT-based binding assay was conducted according to our previous procedure developed for other GPCRs [31–33]. Briefly, HEK293T cells were transfected with an sLgBiT-fused GPR25 expression construct (PB-TRE/sLgBiT-Dr-GPR25, PB-TRE/sLgBiT-Lc-GPR25, or PB-TRE/sLgBiT-Hs-GPR25) with or without the TPST1 and TPST2 coexpression construct pTRE3G-BI/TPST1:TPST2 using the transfection reagent Lipo8000 (Beyotime Technology). Next day, the transfected cells were trypsinized, seeded into white opaque 96-well plates, and cultured in induction medium (complete DMEM plus 20 ng/mL of Dox) at 37 °C for ∼24 h to ∼90% confluence. To start the binding assay, the induction medium was removed and pre-warmed binding solution (serum-free DMEM plus 1% BSA) containing indicated concentrations of tracer (6×His-SmBiT-CXCL17 or 6×His-SmBiT-CXCL17-C26) with or without 1.0 μM of WT or [desC3]CXCL17 was added (50 μL/well). To measure cell surface expression level of the sLgBiT-fused receptors, binding solution containing 80 nM of synthetic HiBiT was added (50 μL/well). After incubation at room temperature for ∼1 h, diluted NanoLuc substrate (30-fold dilution in the binding solution) was added (10 μL/well) and bioluminescence was immediately measured on a SpectraMax iD3 plate reader (Molecular Devices). The measured bioluminescence data were expressed as mean ± SD (*n* = 3) and plotted using the SigmaPlot 10.0 software (SYSTAT software).

## Results

### Human CXCL17 efficiently activates fish GPR25 orthologs in β-arrestin recruitment assays

To investigate potential cross-species activity of human CXCL17, we examined two representative fish GPR25 orthologs: Dr-GPR25 from the ray-finned fish *Danio rerio* (zebrafish) and Lc-GPR25 from the lobe-finned fish *Latimeria chalumnae* (coelacanth) (Table S1). Both orthologs share considerable sequence similarity with human GPR25 (Hs-GPR25), indicating evolutionary conservation and potential functional relevance (Fig. S2).

We employed a NanoBiT-based β-arrestin recruitment assay previously validated for Hs-GPR25 activation [29]. In this assay, the fish GPR25 orthologs were C-terminally fused to an inactive large NanoLuc fragment (LgBiT) and coexpressed with N-terminally SmBiT-tagged human β-arrestin 2 (SmBiT-ARRB2) in HEK293T cells. Upon ligand-induced receptor activation, the proximity of LgBiT and SmBiT restores luciferase activity. Given the high sequence similarity among β-arrestin orthologs from human, zebrafish, and coelacanth (Fig. S3), cross-recognition of human β-arrestin 2 by fish GPR25 orthologs is possible.

Following induction of Dr-GPR25-LgBiT and SmBiT-ARRB2 coexpression in HEK293T cells, basal bioluminescence was observed after NanoLuc substrate addition (Fig. 1A). Upon addition of wild-type (WT) human CXCL17, a rapid and dose-dependent increase in bioluminescence was detected, reaching a maximal effect (∼5-fold over background) with an EC□□ of approximately 60 nM (Fig. 1A). In contrast, [desC3]CXCL17, a CXCL17 mutant lacking three C-terminal residues, failed to elicit a response even at 1.0 μM (Fig. 1B), suggesting the C-terminal region is essential for receptor activation, consistent with previous findings for Hs-GPR25 [29].

**Fig. 1.**
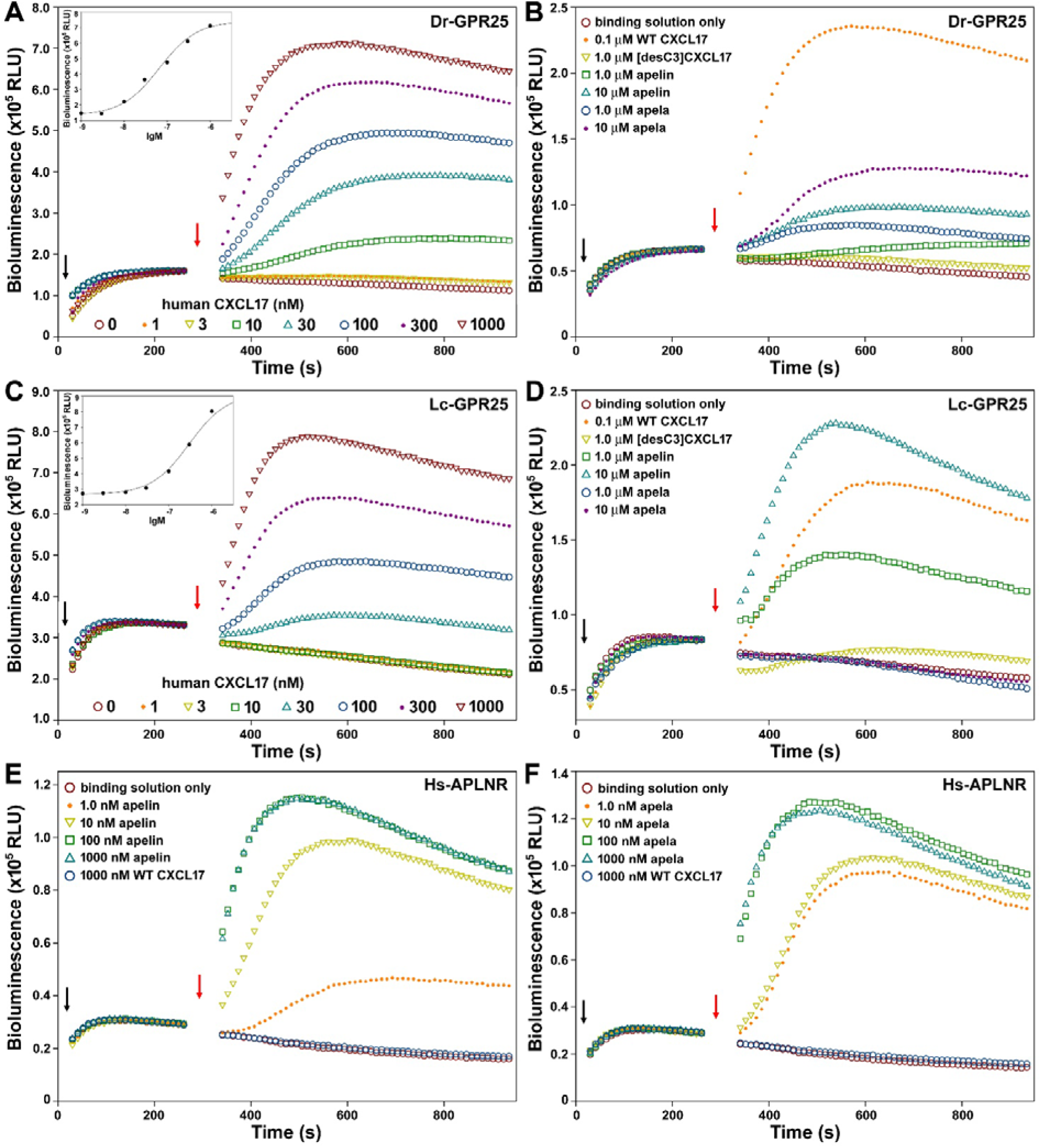
NanoBiT-based β-arrestin recruitment assays for fish GPR25 orthologs and human apelin receptor. (**A**) A typical bioluminescence change after sequential addition of NanoLuc substrate and different concentrations of WT human CXCL17 to living HEK293T cells coexpressing Dr-GPR25-LgBiT and SmBiT-ARRB2. **Inner panel**, dose curve of WT human CXCL17 activating Dr-GPR25 assayed as β-arrestin recruitment. (**B**) A typical bioluminescence change after sequential addition of NanoLuc substrate and [desC3]CXCL17, human apelin-13, or human apela-21 to living HEK293T cells coexpressing Dr-GPR25-LgBiT and SmBiT-ARRB2. (**C**) A typical bioluminescence change after sequential addition of NanoLuc substrate and different concentration of WT human CXCL17 to living HEK293T cells coexpressing Lc-GPR25-LgBiT and SmBiT-ARRB2. **Inner panel**, dose curve of WT human CXCL17 activating Lc-GPR25 assayed as β-arrestin recruitment. (**D**) A typical bioluminescence change after sequential addition of NanoLuc substrate and [desC3]CXCL17, human apelin-13, or human apela-21 to living HEK293T cells coexpressing Lc-GPR25-LgBiT and SmBiT-ARRB2. (**E**) A typical bioluminescence change after sequential addition of NanoLuc substrate and human apelin-13 or WT human CXCL17 to living HEK293T cells coexpressing Hs-APLNR-LgBiT and SmBiT-ARRB2. (**F**) A typical bioluminescence change after sequential addition of NanoLuc substrate and human apela-21 or WT human CXCL17 to living HEK293T cells coexpressing Hs-APLNR-LgBiT and SmBiT-ARRB2. In panel A-F, black arrows indicate the addition of NanoLuc substrate, and red arrows indicate the addition of peptide. The results in these panels are representative of at least two independent experiments.

Synthetic human apelin-13 and apela-21 exhibited minimal activity toward Dr-GPR25 even at high concentrations (Fig. 1B), corroborating earlier reports that they are weak agonists of some non-mammalian GPR25 orthologs [30]. Considering high sequence similarity across species (Fig. S4), apelin and apela are unlikely endogenous agonists for GPR25 in fish species.

Similarly, WT human CXCL17 activated Lc-GPR25 in a dose-dependent manner, with a maximal effect of ∼3-fold over background and an EC□□ of ∼300 nM (Fig. 1C). [desC3]CXCL17 again showed no activation (Fig. 1D). Apelin-13 induced weak activation, whereas apela-21 was inactive toward Lc-GPR25 (Fig. 1D). To verify the integrity of the synthetic apelin and apela, we assessed their activity on their native receptor, human apelin receptor (Hs-APLNR), using the same NanoBiT-based β-arrestin recruitment assay. Both ligands effectively activated Hs-APLNR at concentrations as low as 1.0 nM (Fig. 1E, F), confirming their high activity and ruling out quality issues.

Furthermore, 1.0 μM of WT CXCL17 did not activate Hs-APLNR (Fig. 1E, F), indicating no cross-reactivity between CXCL17 and the apelin receptor. Collectively, these results demonstrated that WT human CXCL17 can efficiently activate both Dr-GPR25 and Lc-GPR25 through its conserved C-terminal domain, suggesting a shared activation mechanism across vertebrate GPR25 orthologs.

### Human CXCL17 efficiently induces chemotactic movement of transfected HEK293T cells expressing fish GPR25 ortholog

Given the chemoattractant role of CXCL17, we evaluated whether it could promote cell migration via fish GPR25 orthologs. Previous work showed that CXCL17 does not induce chemotaxis in untransfected HEK293T cells, indicating the absence of endogenous CXCL17 receptors [29]. Upon transient transfection and induction of Dr-GPR25 or Lc-GPR25 expression, HEK293T cells exhibited dose-dependent chemotactic migration in response to WT human CXCL17 in a transwell assay (Fig. 2). Notably, 100 nM CXCL17 significantly increased cell migration, while [desC3]CXCL17 showed no detectable effect even at 1000 nM. These findings align with the β-arrestin recruitment data and confirm that CXCL17 can induce chemotaxis via fish GPR25 activation.

**Fig. 2.**
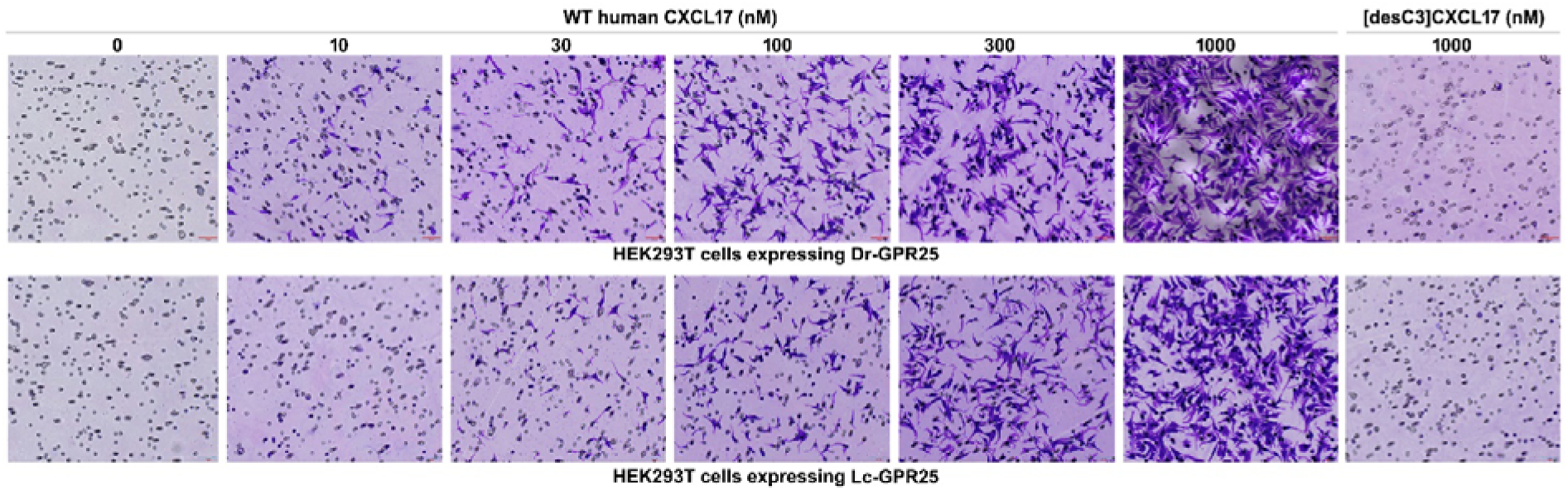
Transwell chemotaxis assay for HEK293T cells expressing fish GPR25 orthologs. Transfected HEK293T cells expressing Dr-GPR25 or Lc-GPR25 were seeded into the permeable membrane-coated inserts and induced by chemotactic solution in the lower chamber. After the assay, cells on the upper face of the permeable membrane were wiped off, and cells on the lower face of the permeable membrane were fixed, stained, and observed under a microscope. The results are representative of two independent experiments.

### Human CXCL17 directly binds to fish GPR25 orthologs in NanoBiT-based binding assays

Following the demonstration that human CXCL17 efficiently activates Dr-GPR25 and Lc-GPR25 and induces chemotactic cell movement, we sought to detect direct binding between human CXCL17 and fish GPR25 orthologs using a NanoBiT-based binding assay, which has been validated with other GPCRs [31–34]. This homogeneous ligand-receptor binding assay involves genetically fusing a secretory large NanoLuc fragment (sLgBiT) to the extracellular N-terminus of a GPCR and covalently attaching a low-affinity SmBiT-tag to an appropriate site on its ligand. Upon binding, a proximity effect induces complementation of the ligand-tagged SmBiT with the receptor-fused sLgBiT, restoring NanoLuc luciferase activity if the SmBiT and sLgBiT are correctly oriented. This NanoBiT-based ligand-receptor binding assay offers high specificity, as endogenously expressed receptors, lacking LgBiT fusion, do not interfere.

To bind to sLgBiT-fused Dr-GPR25 or Lc-GPR25, we initially tested an N-terminally 6×His-SmBiT-fused human CXCL17 (Fig. S1). This tracer significantly increased bioluminescence in the β-arrestin recruitment assay for both Dr-GPR25 and Lc-GPR25 (Fig. 3A,B), suggesting its ability to bind and activate both fish GPR25 orthologs. However, no specific binding was detected when this tracer was added to HEK293T cells overexpressing the N-terminally sLgBiT-fused Dr-GPR25 or Lc-GPR25 (data not shown). This implied that the ligand-fused SmBiT might not complement with the receptor-fused sLgBiT after ligand-receptor binding, likely due to incorrect orientations.

**Fig. 3.**
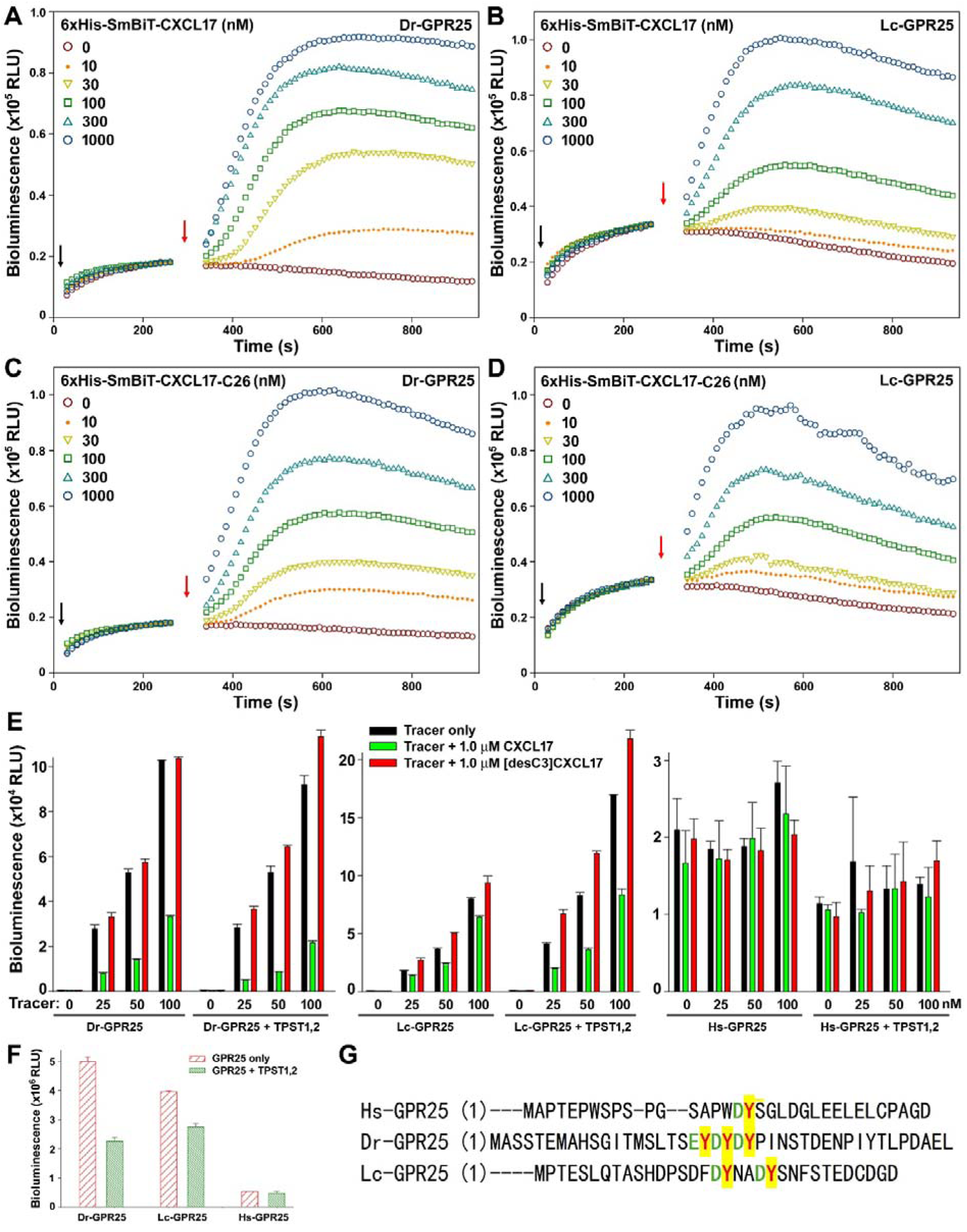
NanoBiT-based ligand-receptor binding assays for fish GPR25 orthologs. (**A,B**) Activity of 6×His-SmBiT-CXCL17 on Dr-GPR25 (A) and Lc-GPR25 (B) measured via NanoBiT-based β-arrestin recruitment assay. (**C,D**) Activity of 6×His-SmBiT-CXCL17-C26 on Dr-GPR25 (C) and Lc-GPR25 (D) measured via NanoBiT-based β-arrestin recruitment assay. In panel C-D, black arrows indicate the addition of NanoLuc substrate, and red arrows indicate the addition of peptide. (**E**) Binding of 6×His-SmBiT-CXCL17-C26 with sLgBiT-fused GPR25 orthologs under absence or presence of 1.0 μM of WT CXCL17 or [desC3]CXCL17. The N-terminally sLgBiT-fused GPR25 ortholog was either expressed alone or coexpressed with human TPST1 and TPST2 in transfected HEK293T cells, and living cells were used for the homogenous binding assay. The measured bioluminescence data are expressed as mean ± SD (*n* = 3) and plotted using the SigmaPlot10.0 software. (**F**) Measurement of cell surface expression level of the sLgBiT-fused GPR25 orthologs in the transfected HEK293T cells via synthetic HiBiT peptide. The same batch of transfected cells in panel E were used to measure cell surface expression level. The measured bioluminescence data are expressed as mean ± SD (*n* = 3) and plotted using the SigmaPlot10.0 software. The results in panel A-F are representative of at least two independent experiments. (**G**) Amino acid sequence of the extracellular N-terminal fragment of GPR25 orthologs. The potentially sulfated Tyr residues are shown in red and highlighted in yellow, and their neighbored negatively charged residues are shown in green.

A previous study indicated that a C-terminal fragment of CXCL17, comprising 26 amino acids (CXCL17-C26), retained considerable activity [28]. Therefore, we designed a 6×His-SmBiT-CXCL17-C26 tracer and overexpressed it as an intein fusion protein in *E. coli* due to its small size (Fig. S1). After renaturation and self-cleavage of the purified intein-fused precursor, 6×His-SmBiT-CXCL17-C26 was released and displayed high activity on both Dr-GPR25 and Lc-GPR25 in the β-arrestin recruitment assay (Fig. 3C,D). This suggested that this shortened tracer could bind to and activate both fish GPR25 orthologs.

Upon adding the 6×His-SmBiT-CXCL17-C26 tracer to living HEK293T cells overexpressing sLgBiT-fused Dr-GPR25 or Lc-GPR25, a significant increase in bioluminescence was observed (Fig. 3E). This increase in bioluminescence could be inhibited by 1.0 μM of WT human CXCL17 but not by 1.0 μM of [desC3]CXCL17 (Fig. 3E). These findings indicate that this shortened tracer can specifically bind to both fish GPR25 orthologs. Some chemokine receptors undergo tyrosine sulfation at their extracellular N-terminal fragment [35,36]. Two tyrosylprotein sulfotransferases, TPST1 and TPST2 (Table S1), are responsible for this posttranslational modification in vertebrates. As shown in Fig. 3G, GPR25 orthologs from human, zebrafish, and coelacanth contain 1-3 potentially sulfated tyrosine residues adjacent to negatively charged residues. When sLgBiT-fused fish GPR25 orthologs were coexpressed with human TPST1 and TPST2, their specific binding with the tracer (bioluminescence without competition minus bioluminescence competed with 1.0 μM of WT CXCL17) significantly increased, particularly for Lc-GPR25 (Fig. 3E). This suggests that tyrosine sulfation enhances the binding of the fish GPR25 orthologs with the tracer.

No specific binding was detected for the 6×His-SmBiT-CXCL17-C26 tracer towards sLgBiT-fused Hs-GPR25 (Fig. 3E). This is likely attributable to the low cell surface expression of human GPR25 in transfected HEK293T cells, as sLgBiT-fused Hs-GPR25 exhibited at least 5-fold lower bioluminescence than the fish orthologs after incubation with the synthetic membrane-impermeable HiBiT peptide (Fig. 3F). Additionally, sLgBiT-fused Hs-GPR25 displayed higher background bioluminescence (without tracer) compared to the fish GPR25 orthologs (Fig. 3E), implying that the N-terminal fragment of human GPR25 might weakly complement with the fused sLgBiT.

## Discussion

This study reveals that human CXCL17 effectively binds to and activates two GPR25 orthologs found in fish, despite the evolutionary divergence of humans and fishes approximately 400 million years ago. These fish GPR25 orthologs exhibit considerably higher cell surface expression in transfected HEK293T cells compared to human GPR25. Consequently, the β-arrestin recruitment assay for the fish GPR25 orthologs is significantly more straightforward than for human GPR25. The maximal effect (E_max_) observed for the fish GPR25 orthologs was at least 3-fold greater than background levels after treatment with 1.0 μM of CXCL17. In contrast, human GPR25, even under optimized conditions, showed at most a one-fold increase above background in our previous study [29]. Therefore, the assays utilizing fish GPR25 orthologs provided robust evidence for CXCL17’s ability to activate GPR25. Given the high efficacy of human CXCL17 with fish GPR25 orthologs, it is reasonable to infer a similarly high activity toward human GPR25, notwithstanding the challenges posed by its low cell surface expression in the β-arrestin recruitment assay, as previously documented [29]. The crucial role of the C-terminal three residues of human CXCL17 in activating both human and fish GPR25 orthologs suggests a conserved mechanism of GPR25 activation across species.

GPR25 orthologs are broadly distributed across vertebrates, from fishes to mammals, and their high evolutionary conservation implies vital functions in all vertebrates. However, CXCL17 orthologs have, to date, only been identified in mammals based on published sequence data. Previous research and the current study indicate that the highly conserved peptide hormones apelin and apela are unlikely to serve as endogenous agonists for non-mammalian GPR25 orthologs, given their notably low activity. Since human CXCL17 maintains high activity toward fish GPR25 orthologs, we speculate that CXCL17 orthologs may indeed exist in non-mammalian vertebrates, even if they have yet to be identified. The modest amino acid sequence similarity among known mammalian CXCL17 orthologs suggests that non-mammalian CXCL17 orthologs might be sufficiently divergent from their mammalian counterparts to preclude their recognition via standard amino acid sequence BLAST searches [29]. Future investigations should aim to test this hypothesis by exploring potential non-mammalian CXCL17 candidates within existing databases and validating their function through established assays, such as the NanoBiT-based β-arrestin recruitment assay.

In summary, our findings reveal that the CXCL17–GPR25 interaction may be conserved across vertebrates, despite the apparent absence of identifiable CXCL17 orthologs outside mammals. This work provides a foundation for exploring the evolutionary conservation and physiological relevance of this chemokine–receptor pair and highlights the value of fish GPR25 orthologs as tractable models for studying orphan GPCRs.

## Abbreviations

BSA: bovine serum albumin
CXCL17: C-X-C motif chemokine ligand 17
DMEM: Dulbecco’s modified Eagle medium
Dox: doxycycline
GPCR: G protein-coupled receptor
GPR25: G protein-coupled receptor 25
HEK: human embryonic kidney
HiBiT: a high-affinity complementation tag for NanoBiT
HPLC: high performance liquid chromatography
LgBiT: a large NanoLuc fragment for NanoBiT
NanoBiT: NanoLuc Binary Technology
PCR: polymerase chain reaction
SD: standard deviation
sLgBiT: secretory LgBiT
SmBiT: a low-affinity complementation tag for NanoBiT
WT: wild-type.

## Author Contributions

WFH, JJW, JY and JJY performed the experiments; YLL and ZGX analyzed the data; ZYG planned the experiments and wrote the paper.

## Acknowledgments

This work was supported by grants from the National Natural Science Foundation of China (31971193, 31470767).

