## Supplemental Tables S1-S2 and Figures S1-S4 for "Human CXCL17 Activates and Binds to Fish GPR25 Orthologs"

### Contents:

**Table S1.** Information about the proteins used in this study.

**Table S2.** Information for generation of the expression constructs for fish GPR25 orthologs.

**Fig. S1.** The nucleotide and amino acid sequence of the SmBiT-based CXCL17 tracers.

**Fig. S2.** Amino acid sequence alignment of GPR25 orthologs from human, zebrafish, and coelacanth.

**Fig. S3.** Amino acid sequence alignment of β-arrestin orthologs from human, zebrafish, and coelacanth.

**Fig. S4.** Amino acid sequence alignment of apelin orthologs and apela orthologs from human, zebrafish, and coelacanth.

**Table S1.** Information about the proteins used in this study. The information is retrieved from gene database of NCBI (https://www.ncbi.nlm.nih.gov/gene).

| Class | Name in this study | Species | Gene ID | mRNA ID | Protein ID | Amino acid sequence |
|---|---|---|---|---|---|---|
| GPR25 orthologs | Hs-GPR25 | *Homo sapiens* (human) | 2848 | NM_005298 | NP_005289 | MAPTEPWSPSPGSAPWDYSGLDGLEELELCPAGDLP<br>YGYVYIPALYLAAFAVGLLGNAFVVWLLAGRRGPRR<br>LVDTFVLHLAAADLGFVLTLPLWAAAAALGGRWPFG<br>DGLCKLSSFALAGTRCAGALLLAGMSVDRYLAVVKL<br>LEARPLRTPRCALASCCGVWAVALLAGLPSLVYRGLQ<br>PLPGGQDSQCGEEPSHAFQGLSLLLLLLTFVLPLVVTL<br>FCYCRISRRLRRPPHVGRARRNSLRIIFAIESTFVGSWL<br>PFSALRAVFHLARLGALPLPCPLLLALRWGLTIATCLA<br>FVNSCANPLIYLLLDRSFRARALDGACGRTGRLARRI<br>SSASSLSRDDSSVFRCRAQAANTASASW |
| | Dr-GPR25 | *Danio rerio* (zebrafish) | 795188 | XM_073916757 | XP_073772858 | MASSTEMAHSGITMSLTSEYDYDYPINSTDENPIYTLP<br>DAELLPMSNIYIPVLYIIMFLTGSLGNLFVIVVIGKRRK<br>KSGRLVDTFVLNLALADLVFVLTLPMWAISTRYDEW<br>PFGEVLCKISSFIIAVNRFSNIFFLTCMSVDRYLAVVRL<br>MDSRFLRSSNCAQITCGIVWVVSFFLGSPSLAYRHLIN<br>NSVCSEDSKSSFVQGMNLLTILLTFLLPVLILGLCYGSI<br>LVNLRRHCHNPANTRTDARRRHSVKIVFAIISAFLISW<br>LPFNCFKAIHVALLIINGDLNEDTYVVIHRGLMLSCCL<br>AFLNSCVNPAIYFFLDQHFRRRASMLCLSCLSQNDQA<br>HQSYITSNSYSNGTSETCSGNTSTRGRLFSLTQKA |
| | Lc-GPR25 | *Latimeria chalumnae* (coelacanth) | 102365624 | XM_005988473 | XP_005988535 | MPTESLQTASHDPSDFDYNADYSNFSTEDCDGDLPY<br>AKIYIPIFYFVIFFTGLFGNVFVIAAMTLKQTTKRLVDI<br>FVINLAVADLVFVFTLPLWSVSAAFDDQWLFGGVLC<br>KLSSYVIAVNRYSSIFFMTGMSVDRYMAVVKLLDSKF<br>IRTRRCILITCTIIWIISLVMGIPSLVYRDLSTQDSEHTY<br>CIEDQDSIIFKGISLASLFLAFVLPVIIILFCYCSISARLY<br>SHFHHANRYDQKRKKTLKIIFTIITAFVCSWLPFNTFK<br>TLYLLFSFQGKMPPCRVGILRQGLTITACFAFLSSCVN<br>PIIYTFLDNHFRKRAHRLLVKALGRYTERRNSFGESW<br>ASETSSTFVSRIRANSVKELQNMNKTQQNTIPT |
| β-arrestin (ARRB) orthologs | Hs-ARRB2 | *Homo sapiens* (human) | 409 | NM_004313 | NP_004304 | MGEKPGTRVFKKSSPNCKLTVYLGKRDFVDHLDKV<br>DPVDGVVLVDPDYLKDRKVFVTLTCAFRYGREDLDV<br>LGLSFRKDLFIATYQAFPPVPNPPRPPTRLQDRLLRKL<br>GQHAHPFFFTIPQNLPCSVTLQPGPEDTGKACGVDFE<br>IRAFCAKSLEEKSHKRNSVRLVIRKVQFAPEKPGPQPS<br>AETTRHFLMSDRSLHLEASLDKELYYHGEPLNVNVH<br>VTNNSTKTVKKIKVSVRQYADICLFSTAQYKCPVAQL<br>EQDDQVSPSSTFCKVYTITPLLSDNREKRGLALDGKL<br>KHEDTNLASSTIVKEGANKEVLGILVSYRVKVKLVVS<br>RGGDVSVELPFVLMHPKPHDHIPLPRPQSAAPETDVP<br>VDTNLIEFDTNYATDDDIVFEDFARLRLKGMKDDDY<br>DDQLC |
| | Dr-ARRB2A | *Danio rerio* (zebrafish) | 321378 | NM_214681 | NP_999846 | MGDKAGTRVFKKSSPNCKVTVYLGKRDFVDHLDHV<br>DPVDGVILVDPEYLKDRKVFVTLTCAFRYGREDLDV<br>LGLSFRKDLYIFTFQAYPPIPEESKPHSRLQERLLKKL<br>GQNAYPFHFSIPQNLPCSVTLQPGPEDTGKACGVDFE<br>IRAFCAKSMEEKNHKRNSVRLVIRKAQYAPEKPGPQP<br>MVETTRSFLMSDRSLHLEASLDKELYYHGEPISVNVH<br>VTNNSTKTVKRVKISVRQYADICLFSTAQYKCPVAQI<br>EADDQVASSSTFCKVYTLTPTLNNNREKRGPALDGK<br>LKHEDTNLASSTIVKDVSNKEVLGVLVSYRVKVKLV |

| | | | | | | |
|---|---|---|---|---|---|---|
| | | | | | | VSRGGDVSVELPFVLMHPKPSESSLSHSTSAVPMLDPPIDTNLIEFDTNSLIPDDDIVFEDFARLRLKGVIDKEEDC |
| | Dr-ARRB2B | *Danio rerio* (zebrafish) | 394099 | NM_201124 | NP_957418 | MGDKAGTRVFKKSSPNCKLTVYLGKRDFVDHLDHVDPVDGVLLIDPEYLKDRKVFVTLTCAFRYGREDLDVLGLSFRKDLFISSFQAYPPLPDERKPLSRLQERLLKKLGQNAYPFNFTIPQNLPCSVTLQPGPEDTGKACGVDFEVRAFCAKTVDEKTHKRNSVRLVIRKVQYAPEKPGPQPMVETTRSFLMSDRSLHLEASLDKELYYHGEPISVNVHVTNNSTKTVKRVKISVRQYADICLFSTAQYKCPVAQVEADDQVSSSSTFCKVYTLTPTLSNNREKRGLALDGQLKHEDTNLASSTIVKDVSNKEVLGILVSYRVKVKLVVSRGGDVSVELPFVLMHPKPSEQPNSRPQSAVPETDVPVDANLIEFETNNFSQDDDFVFEDFARLRLKGMKDEEDDHFC |
| | Lc-ARRB1 | *Latimeria chalumnae* (coelacanth) | 102360512 | XM_064556990 | XP_064413060 | MDSFSITWRVIGSSFSTDRASLKDSAALSVTVWQIVRVFKKASPNGKLTVYLGKRDFVDHVDFVDPVDGVVLVDPEYLKERKVFVTLTCAFRYGREDLDVLGLTFRKDLFVANVQAFPPVPEEKKPLTRLQERLIKKLGEHAYPFTFEIPPNLPCSVTLQPGPEDTGKACGVDFEVKTFCAENLEEKIHKRNSVRLVIRKVQYAPEKPGPQPMAETTRQFLMSDKPLHLEASLDKEIYYHGEPISVNVHVTNNTNKTVKKIKISVRQYADICLFNTAQYKCPVAVEEADDVVAPSSTFCKVYTLVPFLANNREKRGLALDGKLKHEDTNLASSTLLRDGANKEILGIIVSYKVKVKLVVSRGGILGDLASSDVAVELPFTLMHPKPKEEALYREVPESEAPIDTNLIEFDTNDDDIVFEDFARQRLKGMKDDKDDDDDVASSPQLNDR |
| Apelin orthologs | Hs-apelin | *Homo sapiens* (human) | 8862 | NM_017413 | NP_059109 | MNLRLCVQALLLLWLSLTAVCGGSLMPLPDGNGLEDGNVRHLVQPRGSRNGPGPWQGGRRKFRRQRPRLSHKGPMPF |
| | Dr-apelin | *Danio rerio* (zebrafish) | 798375 | NM_001166124 | NP_001159596 | MNVKILTLVIVLVVSLLCSASAGPMASTEHSKEIEEVGSMRTPLRQNPARAGRSQRPAGWRRRRPRPRLSHKGPMPF |
| | Lc-apelin | *Latimeria chalumnae* (coelacanth) | 106702803 | XM_014486099 | XP_014341585 | MNCKSLLFGVLFLFLTLTSVWGGPLAIPLEGSDLEEGSLKNLVQQSIVRNNSGHRQSGWRRYRRPRPRLSHKGPMPF |
| Apela orthologs | Hs-apela | *Homo sapiens* (human) | 100506013 | NM_001297550 | NP_001284479 | MRFQQFLFAFFIFIMSLLLISGQRPVNLTMRRKLRKHNCLQRRCMPLHSRVPFP |
| | Dr-apela | *Danio rerio* (zebrafish) | 100536023 | NM_001297547 | NP_001284476 | MRFFHPLYLLLLLLTVLVLISADKHGTKHDFLNLRRKYRRHNCPKKRCLPLHSRVPFP |
| | Lc-apela | *Latimeria chalumnae* (coelacanth) | 102356079 | XM_014497274 | XP_014352760 | MRFQQLFQILFLLLLSLLLIHGDGPANLSSRRRKLRHKCPHRRCMALHSRVPFP |
| Apelin receptor | Hs-APLNR | *Homo sapiens* (human) | 187 | NM_005161 | NP_005152 | MEEGGDFDNYYGADNQSECEYTDWKSSGALIPAIYMLVFLLGTTGNGLVLWTVFRSSREKRRSADIFIASLAVADLTFVVTLPLWATYTYRDYDWPFGTFFCKLSSYLIFVNMYASVFCLTGLSFDRYLAIVRPVANARLRLRVSGAVATAVLWVLAALLAMPVMVLRTTGDLENTTKVQCYMDYSMVATVSSEWAWEVGLGVSSTTVGFVVPFTIMLTCYFFIAQTIAGHFRKERIEGLRKRRRLLSIIVVLVVTFALCWMPYHLVKTLYMLGSLLHWPCDFDLFLMNIFPYCTCISYVNSCLNPFLYAFFDPRFRQACTSMLCCGQSRCAGTSHSSSGEKSASYSSGHSQGPGPNMGKGGEQMHEKSIPYSQETLVVD |

| | | | | | | |
|---|---|---|---|---|---|---|
| TPST | TPST1 | *Homo sapiens* (human) | 8460 | NM_003596 | NP_003587 | MVGKLKQNLLLACLVISSVTVFYLGQHAMECHHRIEERSQPVKLESTRTTVRTGLDLKANKTFAYHKDMPLIFIGGVPRSGTTLMRAMLDAHPDIRCGEETRVIPRILALKQMWSRSSKEKIRLDEAGVTDEVLDSAMQAFLLEIIVKHGEPAPYLCNKDPFALKSLTYLSRLFPNAKFLLMVRDGRASVHSMISRKVTIAGFDLNSYRDCLTKWNRAIETMYNQCMEVGYKKCMLVHYEQLVLHPERWMRTLLKFLQIPWNHSVLHHEEMIGKAGGVSLSKVERSTDQVIKPVNVGALSKWVGKIPPDVLQDMAVIAPMLAKLGYDPYANPPNYGKPDPKIIENTRRVYKGEFQLPDFLKEKPQTEQVE |
| | TPST2 | *Homo sapiens* (human) | 8459 | NM_001008566 | NP_001008566 | MRLSVRRVLLAAGCALVLVLAVQLGQQVLECRAVLAGLRSPRGAMRPEQEELVMVGTNHVEYRYGKAMPLIFVGGVPRSGTTLMRAMLDAHPEVRCGEETRIIPRVLAMRQAWSKSGREKLRLDEAGVTDEVLDAAMQAFILEVIAKHGEPARVLCNKDPFTLKSSVYLSRLFPNSKFLLMVRDGRASVHSMITRKVTIAGFDLSSYRDCLTKWNKAIEVMYAQCMEVGKEKCLPVYYEQLVLHPRRSLKLILDFLGIAWSDAVLHHEDLIGKPGGVSLSKIERSTDQVIKPVNLEALSKWTGHIPGDVVRDMAQIAPMLAQLGYDPYANPPNYGNPDPFVINNTQRVLKGDYKTPANLKGYFQVNQNSTSSHLGSS |

**Table S2.** Information for generation of the expression constructs for fish GPR25 orthologs.

| Expression constructs | Vectors for cloning | Restriction enzymes cleaving the vector | Primers for PCR amplification (5' to 3') | Template for PCR amplification | Approach for construct generation |
|---|---|---|---|---|---|
| pTRE3G-BI/ Dr-GPR25-LgBiT: SmBiT-ARRB2 | pTRE3G-BI/ MRGPRX2-LgBiT: SmBiT-ARRB2 | NheI; AgeI (removing MRGPRX2) | Forward: CGG GGA GAC CCA AGC TGG ATG GCA AGC AGC ACA GAG ATG<br>Reverse: CCC TCC GCC GGT ACC GAC CGG TGG TGC CTT TTG AGT AAG TGA GAA AA | Zebrafish genomic DNA | Gibson assembly |
| PB-TRE/sLgBiT-Dr-GPR25 | PB-TRE/ sLgBiT-GPR182 | KpnI; PmeI (removing GPR182) | Forward: GGT GGC AGC GGC GGT GGT ACC GCA AGC AGC ACA GAG ATG GCA<br>Reverse: GAG GCT GAT CAG CGG GT TTA TGC CTT TTG AGT AAG TGA GAA | pTRE3G-BI/ Dr-GPR25-LgBiT: SmBiT-ARRB2 | Gibson assembly |
| PB-TRE/Dr-GPR25 | PB-TRE/ dCas9-VPR | NheI; PmeI (removing dCas9-VPR) | Forward: CCC TCG TAA AGG TCT AGA G ATG GCA AGC AGC ACA GAG AT<br>Reverse: GTT TCA GTT AGC CTC CCC CGT TTA TGC CTT TTG AGT AAG TGA GAA | pTRE3G-BI/ Dr-GPR25-LgBiT: SmBiT-ARRB2 | Gibson assembly |
| pcDNA3.1/Lc-GPR25 | pcDNA3.1 | NheI; NotI | Chemical synthesis of Lc-GPR25 coding region | No | Gibson assembly |
| pTRE3G-BI/ Lc-GPR25-LgBiT: SmBiT-ARRB2 | pTRE3G-BI/ MRGPRX2-LgBiT: SmBiT-ARRB2 | NheI; AgeI (removing MRGPRX2) | Forward: CCG TCA GAT CGC CTG GAG AAT TCG GGG AGA CCC AAG CTG GCT AGC<br>Reverse: GCT AGA CCC TCC GCC GGT ACC GAC CGG TGG AGT AGG TAT TGT GTT CTG CTG | pcDNA3.1/ Lc-GPR25 | Gibson assembly |
| PB-TRE/sLgBiT-Lc-GPR25 | PB-TRE/ sLgBiT-GPR182 | KpnI; PmeI (removing GPR182) | Forward: GGT GGC AGC GGC GGT GGT ACC CCA ACT GAA AGT TTA CAA A<br>Reverse: GAG GCT GAT CAG CGG GT TTA AGT AGG TAT TGT GTT CT | pcDNA3.1/ Lc-GPR25 | Gibson assembly |
| PB-TRE/Lc-GPR25 | PB-TRE/ dCas9-VPR | NheI; PmeI (removing dCas9-VPR) | Forward: TTC CTA CCC TCG TAA AGG TCT AGA G CTC ACT ATA GGG AGA CCC AAG CT<br>Reverse: GTT TCA GTT AGC CTC CCC CGT TT C ACA GTC GAG GCT GAT CAG CGG | pcDNA3.1/ Lc-GPR25 | Gibson assembly |

PB-TRE/sLgBiT-GPR182 was generated in our laboratory based on PB-TRE/dCas9-VPR (Addgene cat#: 63800) via removal of the dCas9-VPR fragment by NheI and PmeI cleavage and then ligation of sLgBiT-GPR182 fragment, unpublished data.

pTRE3G-BI/GPR83-LgBiT:SmBiT-ARRB2 was generated in our previous study.

For oligo primers, the sequence pairing with vector is highlighted in yellow, the sequence pairing with PCR template is underlined.

#### 6xHis-SmBiT-CXCL17

```
1    ATG CAT CAC CAT CAC CAC CAT GGT GGC GTG ACC GGC TAC CGT CTG TTT GAA GAA ATT CTG GGC GGC AGC GGT GGT
     TAC GTA GTG GTA GTG GTG GTA CCA CCG CAC TGG CCG ATG GCA GAC AAA CTT CTT TAA GAC CCG CCG TCG CCA CCA
      M   H   H   H   H   H   H   G   G   V   T   G   Y   R   L   F   E   E   I   L   G   G   S   G   G

76   GGC AGC AGT CTG AAT CCA GGT GTC GCA CGT GGC CAC CGC GAC CGT GGC CAG GCT TCT CGC CGT TGG CTC CAG GAA
     CCG TCG TCA GAC TTA GGT CCA CAG CGT GCA CCG GTG GCG CTG GCA CCG GTC CGA AGA GCG GCA ACC GAG GTC CTT
      G   S   S   L   N   P   G   V   A   R   G   H   R   D   R   G   Q   A   S   R   R   W   L   Q   E

151  GGT GGC CAA GAA TGT GAG TGC AAA GAT TGG TTC CTG CGT GCC CCG CGC CGT AAA TTC ATG ACC GTG TCC GGT CTG
     CCA CCG GTT CTT ACA CTC ACG TTT CTA ACC AAG GAC GCA CGG GGC GCG GCA TTT AAG TAC TGG CAC AGG CCA GAC
      G   G   Q   E   C   E   C   K   D   W   F   L   R   A   P   R   R   K   F   M   T   V   S   G   L

226  CCG AAG AAA CAG TGC CCT TGT GAT CAT TTC AAG GGC AAT GTT AAG AAA ACG CGC CAC CAA CGT CAT CAC CGT AAG
     GGC TTC TTT GTC ACG GGA ACA CTA GTA AAG TTC CCG TTA CAA TTC TTT TGC GCG GTG GTT GCA GTA GTG GCA TTC
      P   K   K   Q   C   P   C   D   H   F   K   G   N   V   K   K   T   R   H   Q   R   H   H   R   K

301  CCG AAC AAA CAT TCA CGC GCG TGC CAG CAA TTT CTC AAA CAA TGT CAG CTT CGT AGC TTT GCT CTG CCT TTG TAA
     GGC TTG TTT GTA AGT GCG CGC ACG GTC GTT AAA GAG TTT GTT ACA GTC GAA GCA TCG AAA CGA GAC GGA AAC ATT
      P   N   K   H   S   R   A   C   Q   Q   F   L   K   Q   C   Q   L   R   S   F   A   L   P   L   *
```

#### 6xHis-SmBiT-CXCL17-C26-Intein

```
1    CTT TAA GAA GGA GAT ATA ATG CAT CAT CAC CAT CAC CAT GGT GTG ACC GGC TAC CGT CTG TTT GAA GAA ATT CTG
     GAA ATT CTT CCT CTA TAT TAC GTA GTA GTG GTA GTG GTA CCA CAC TGG CCG ATG GCA GAC AAA CTT CTT TAA GAC
                              M   H   H   H   H   H   H   G   V   T   G   Y   R   L   F   E   E   I   L

76   GGC GGC GGT GGT CGT AAG CCG AAC AAA CAT TCA CGC GCG AGC CAG CAA TTT CTC AAA CAA TCT CAG CTT CGT AGC
     CCG CCG CCA CCA GCA TTC GGC TTG TTT GTA AGT GCG CGC TCG GTC GTT AAA GAG TTT GTT AGA GTC GAA GCA TCG
      G   G   G   G   R   K   P   N   K   H   S   R   A   S   Q   Q   F   L   K   Q   S   Q   L   R   S

151  TTT GCT CTG CCT TTG TGT ATT TGC GGT GAT GCT CTG GTG GCC CTG CCG GAA GGC GAA AGC GTT CGT ATC GCA GAC
     AAA CGA GAC GGA AAC ACA TAA ACG CCA CTA CGA GAC CAC CGG GAC GGC CTT CCG CTT TCG CAA GCA TAG CGT CTG
      F   A   L   P   L   C   I   C   G   D   A   L   V   A   L   P   E   G   E   S   V   R   I   A   D

226  ATT GTC CCA GGT GCT CGC CCG AAC AGT GAT AAT GCG ATC GAC CTG AAA GTA CTC GAT CGT CAC GGC AAC CCA GTG
     TAA CAG GGT CCA CGA GCG GGC TTG TCA CTA TTA CGC TAG CTG GAC TTT CAT GAG CTA GCA GTG CCG TTG GGT CAC
      I   V   P   G   A   R   P   N   S   D   N   A   I   D   L   K   V   L   D   R   H   G   N   P   V

301  TTA GCG GAT CGT TTG TTT CAT TCT GGT GAG CAT CCT GTT TAT ACC GTC CGC ACG GTA GAA GGT CTG CGT GTG ACT
     AAT CGC CTA GCA AAC AAA GTA AGA CCA CTC GTA GGA CAA ATA TGG CAG GCG TGC CAT CTT CCA GAC GCA CAC TGA
      L   A   D   R   L   F   H   S   G   E   H   P   V   Y   T   V   R   T   V   E   G   L   R   V   T

376  GGC ACC GCC AAC CAC CCG CTG CTG TGC TTA GTT GAT GTC GCA GGT GTA CCG ACC CTG TTG TGG AAG TTA ATT GAT
     CCG TGG CGG TTG GTG GGC GAC GAC ACG AAT CAA CTA CAG CGT CCA CAT GGC TGG GAC AAC ACC TTC AAT TAA CTA
      G   T   A   N   H   P   L   L   C   L   V   D   V   A   G   V   P   T   L   L   W   K   L   I   D

451  GAG ATC AAA CCA GGC GAC TAC GCT GTG ATT CAG CGT TCC GCG TTC TCA GTT GAT TGT GCT GGC TTT GCC CGC GGT
     CTC TAG TTT GGT CCG CTG ATG CGA CAC TAA GTC GCA AGG CGC AAG AGT CAA CTA ACA CGA CCG AAA CGG GCG CCA
      E   I   K   P   G   D   Y   A   V   I   Q   R   S   A   F   S   V   D   C   A   G   F   A   R   G

526  AAA CCG GAG TTC GCA CCT ACC ACT TAT ACG GTG GGC GTT CCG GGT CTG GTC CGT TTC CTG GAG GCG CAC CAT CGC
     TTT GGC CTC AAG CGT GGA TGG TGA ATA TGC CAC CCG CAA GGC CCA GAC CAG GCA AAG GAC CTC CGC GTG GTA GCG
      K   P   E   F   A   P   T   T   Y   T   V   G   V   P   G   L   V   R   F   L   E   A   H   H   R

601  GAT CCA GAC GCA CAG GCT ATC GCG GAT GAA CTG ACG GAT GGC CGT TTT TAT TAC GCC AAA GTG GCT AGC GTT ACC
     CTA GGT CTG CGT GTC CGA TAG CGC CTA CTT GAC TGC CTA CCG GCA AAA ATA ATG CGG TTT CAC CGA TCG CAA TGG
      D   P   D   A   Q   A   I   A   D   E   L   T   D   G   R   F   Y   Y   A   K   V   A   S   V   T

676  GAC GCG GGT GTG CAA CCG GTC TAT TCG TTA CGC GTT GAT ACG GCA GAC CAC GCG TTC ATC ACG AAC GGC TTC GTT
     CTG CGC CCA CAC GTT GGC CAG ATA AGC AAT GCG CAA CTA TGC CGT CTG GTG CGC AAG TAG TGC TTG CCG AAG CAA
      D   A   G   V   Q   P   V   Y   S   L   R   V   D   T   A   D   H   A   F   I   T   N   G   F   V

751  TCT CAT GCG TAA GCG GCC GCA CTC GAG CAC CAC CAC CAC CAC CAC TGA
     AGA GTA CGC ATT CGC CGG CGT GAG CTC GTG GTG GTG GTG GTG GTG ACT
      S   H   A   *
```

**Fig. S1.** The nucleotide and amino acid sequence of the SmBiT-based CXCL17 tracers. The amino acid sequence of SmBiT is shown in blue, that of intact human CXCL17 or its C26 fragment in red, and that of intein in green. Two Cys residues of human CXCL17 are replaced by a Ser residue in the C26 fragment and highlighted in yellow.

```
Hs-GPR25  (1) ---MAPTEPWSPS-PG----SAPWDYSGLDGLEELELCPAGDLPYGYVYIPALYLAAFAVGLLGNAFVVWLLAGRRGPR-RLVDTFVLHLAAADLGFVLTLPLWAAAAALGGRWPFGDGLCKLSSFALAG
Dr-GPR25  (1) MASSTEMAHSGITMSLTSEYDYDYPINSTDENPIYTLPDAELLPMSNIYIPVLYIIMFLTGSLGNLFVIVVIGKRRKKSGRLVDTFVLNLALADLVFVLTLPMWAISTRY-DEWPFGEVLCKISSFIIAV
Lc-GPR25  (1) ----MPTESLQTASHDP----SDFDYNADYSNFSTEDCDG-DLPYAKIYIPIFYFVIFFTGLFGNVFVIAAMTLKQTTK-RLVDIFVINLAVADLVFVFTLPLWSVSAAFDDQWLFGGVLCKLSSYVIAV

Hs-GPR25  (122) TRCAGALLLAGMSVDRYLAVVKLLEARPLRTPRCALASCCGVWAVALLAGLPSLVYRGLQPLPGGQDSQCGEEPSHAFQGLSLLLLLTFVLPLVVTLFCYCRISRRLRRPPHVG------RARRNSLRI
Dr-GPR25  (130) NRFSNIFFLTCMSVDRYLAVVRLMDSRFLRSSNCAQITCGIVWVVSFFLGSPSLAYRHLINN----SVCSEDSKSSFVQGMNLLTILLTFLLPVLILGLCYGSILVNLRRHCHNPANTRTDARRRHSVKI
Lc-GPR25  (121) NRYSSIFFMTGMSVDRYMAVVKLLDSKFIRTRRCILITCTIIWIISLVMGIPSLVYRDLSTQDSEHTYCIEDQDSIIFKGISLASLFLAFVLPVIIILFCYCSISARLYSHFHHAN--RYDQKRKKTLKI

Hs-GPR25  (246) IFAIESTFVGSWLPFSALRAVFHLARLGALPLPCPLLLALRWGLTIATCLAFVNSCANPLIYLLLDRSFRARALDGACGRTGRLARRISSASS--LSRDDSSVFRCRAQAANTASASW-----------
Dr-GPR25  (256) VFAIISAFLISWLPFNCFKAIHVALLIINGDLNEDTYVVIHRGLMLSCCLAFLNSCVNPAIYFFLDQHFRRRASMLCLSCLSQNDQAHQSYITSNSYSNGTSETCSGNTSTRGRLFSLTQKA-------
Lc-GPR25  (249) IFTIITAFVCSWLPFNTFKTLYLLFSFQGKMPPCRVG-ILRQGLTITACFAFLSSCVNPIIYTFLDNHFRKRAHRLLVKALGRYTERRNSFGES-WASETSSTFVSRIRANSVKELQNMNKTQQNTIPT
```

**Fig. S2.** Amino acid sequence alignment of GPR25 orthologs from human, zebrafish, and coelacanth. These GPR25 orthologs are retrieved from the gene database of NCBI (Table S1) and aligned via Vector NTI software.

```
Hs-ARRB2     (1)  -----------------------------MGEKPGTRVFKKSSPNCKLTVYLGKRDFVDHLDKVDPVDGVVLVDPDYLKDRKVFVTLTCAFRYGREDLDVLGLSFRKDLFIATYQ
Dr-ARRB2A    (1)  -----------------------------MGDKAGTRVFKKSSPNCKVTVYLGKRDFVDHLDHVDPVDGVILVDPEYLKDRKVFVTLTCAFRYGREDLDVLGLSFRKDLYIFTFQ
Dr-ARRB2B    (1)  -----------------------------MGDKAGTRVFKKSSPNCKLTVYLGKRDFVDHLDHVDPVDGVLLIDPEYLKDRKVFVTLTCAFRYGREDLDVLGLSFRKDLFISSFQ
Lc-ARRB1     (1)  MDSFSITWRVIGSSFSTDRASLKDSAALSVTVWQIVRVFKKASPNGKLTVYLGKRDFVDHVDFVDPVDGVVLVDPEYLKERKVFVTLTCAFRYGREDLDVLGLTFRKDLFVANVQ

Hs-ARRB2    (87)  AFPPVPNPPRPPTRLQDRLLRKLGQHAHPFFFTIPQNLPCSVTLQPGPEDTGKACGVDFEIRAFCAKSLEEKSHKRNSVRLVIRKVQFAPEKPGPQPSAETTRHFLMSDRSLHLE
Dr-ARRB2A   (87)  AYPPIPEESKPHSRLQERLLKKLGQNAYPFHFSIPQNLPCSVTLQPGPEDTGKACGVDFEIRAFCAKSMEEKNHKRNSVRLVIRKAQYAPEKPGPQPMVETTRSFLMSDRSLHLE
Dr-ARRB2B   (87)  AYPPLPDERKPLSRLQERLLKKLGQNAYPFNFTIPQNLPCSVTLQPGPEDTGKACGVDFEVRAFCAKTVDEKTHKRNSVRLVIRKVQYAPEKPGPQPMVETTRSFLMSDRSLHLE
Lc-ARRB1    (116) AFPPVPEEKKPLTRLQERLIKKLGEHAYPFTFEIPPNLPCSVTLQPGPEDTGKACGVDFEVKTFCAENLEEKIHKRNSVRLVIRKVQYAPEKPGPQPMAETTRQFLMSDKPLHLE

Hs-ARRB2    (202) ASLDKELYYHGEPLNVNVHVTNNSTKTVKKIKVSVRQYADICLFSTAQYKCPVAQLEQDDQVSPSSTFCKVYTITPLLSDNREKRGLALDGKLKHEDTNLASSTIVKEGANKEVL
Dr-ARRB2A   (202) ASLDKELYYHGEPISVNVHVTNNSTKTVKRVKISVRQYADICLFSTAQYKCPVAQIEADDQVASSSTFCKVYTLTPTLNNNREKRGPALDGKLKHEDTNLASSTIVKDVSNKEVL
Dr-ARRB2B   (202) ASLDKELYYHGEPISVNVHVTNNSTKTVKRVKISVRQYADICLFSTAQYKCPVAQVEADDQVSSSSTFCKVYTLTPTLSNNREKRGLALDGQLKHEDTNLASSTIVKDVSNKEVL
Lc-ARRB1    (231) ASLDKEIYYHGEPISVNVHVTNNTNKTVKKIKISVRQYADICLFNTAQYKCPVAVEEADDVVAPSSTFCKVYTLVPFLANNREKRGLALDGKLKHEDTNLASSTLLRDGANKEIL

Hs-ARRB2    (317) GILVSYRVKVKLVVSRG--------GDVSVELPFVLMHPKPHDHIPLPRPQSAAPETDVPVDTNLIEFDTN-YATDDDIVFEDFARLRLKGMKDDDYDDQLC---------
Dr-ARRB2A   (317) GVLVSYRVKVKLVVSRG--------GDVSVELPFVLMHPKPSESSLS-HSTSAVPMLDPPIDTNLIEFDTNSLIPDDDIVFEDFARLRLKGVIDKEEDC-----------
Dr-ARRB2B   (317) GILVSYRVKVKLVVSRG--------GDVSVELPFVLMHPKPSEQPNS-RPQSAVPETDVPVDANLIEFETNNFSQDDDFVFEDFARLRLKGMKDEEDDHFC----------
Lc-ARRB1    (346) GIIVSYKVKVKLVVSRGGILGDLASSDVAVELPFTLMHPKPKEEALY----REVPESEAPIDTNLIEFDTN----DDDIVFEDFARQRLKGMKDDKDDDDDVASSPQLNDR
```

**Fig. S3.** Amino acid sequence alignment of β-arrestin orthologs from human, zebrafish, and coelacanth. These β-arrestin orthologs are retrieved from the gene database of NCBI (Table S1) and aligned via Vector NTI software.

```
Hs-apelin   (1) MNLRLCVQALLLLWLSLTAVCGGSLMPLPDGNGLED-GNVRHLVQPRGSRNGPGPWQGGRRKFRRQRPRLSHKGPMPF
Dr-apelin   (1) MNVKILTLVIVLVVSLLCSASAGPMASTEHSKEIEEVGSMRTPLRQNPARAGRSQRPAGWR-RRRPRPRLSHKGPMPF
Lc-apelin   (1) MNCKSLLFGVLFLFLTLTSVWGGPLAIPLEGSDLEE-GSLKNLVQQSIVRNNSGHRQSGWRRYRRPRPRLSHKGPMPF
```

```
Hs-apela   (1) MRFQQFLFAFFIFIMSLLLISGQRP----VNLTMRRKLRKHNCLQRRCMPLHSRVPFP
Dr-apela   (1) MRFFHPLYLLLLLLTVLVLISADKHGTKHDFLNLRRKYRRHNCPKKRCLPLHSRVPFP
Lc-apela   (1) MRFQQLFQILFLLLLSLLLIHGDGP----ANLSSRRRKLRHKCPHRRCMALHSRVPFP
```

**Fig. S4.** Amino acid sequence alignment of apelin orthologs and apela orthologs from human, zebrafish, and coelacanth. These orthologs are retrieved from the gene database of NCBI (Table S1) and aligned via Vector NTI software. The sequences of human apelin-13 and human apela-21 are underlined.
